# Assessing the accuracy of Approximate Bayesian Computation approaches to infer epidemiological parameters from phylogenies

**DOI:** 10.1101/050211

**Authors:** Emma Saulnier, Olivier Gascuel, Samuel Alizon

## Abstract

Phylodynamics typically rely on likelihood-based methods to infer epidemiological parameters from dated phylogenies. These methods are essentially based on simple epidemiological models because of the difficulty in expressing the likelihood function analytically. Computing this function numerically raises additional challenges, especially for large phylogenies. Here, we use Approximate Bayesian Computation (ABC) to circumvent these problems. ABC is a likelihood-free method of parameter inference, based on simulation and comparison between target data and simulated data, using summary statistics. We simulated target trees under several epidemiological scenarios in order to assess the accuracy of ABC methods for inferring epidemiological parameter such as the basic reproduction number (*R*_0_), the mean duration of infection, and the effective host population size. We designed many summary statistics to capture the information in a phylogeny and its corresponding lineage-through-time plot. We then used the simplest ABC method, called rejection, and its modern derivative complemented with adjustment of the posterior distribution by regression. The availability of machine learning techniques including variable selection, motivated us to compute many summary statistics on the phylogeny. We found that ABC-based inference reaches an accuracy comparable to that of likelihood-based methods for birth-death models and can even outperform existing methods for more refined models and large trees. By re-analysing data from the early stages of the recent Ebola epidemic in Sierra Leone, we also found that ABC provides more realistic estimates than the likelihood-based methods, for some parameters. This work shows that the combination of ABC-based inference using many summary statistics and sophisticated machine learning methods able to perform variable selection is a promising approach to analyse large phylogenies and non-trivial models.

## Introduction

To control epidemics, we must understand their dynamics. Classical analyses typically rely on prevalence or incidence data [1,2], which correspond to the total number of reported cases, and the number of newly reported cases through time, respectively. By combining such data with epidemiological models, one can estimate key parameters, such as the basic reproduction number (*R*_0_), which is the number of secondary cases generated by an infectious individual in a fully susceptible host population. A robust and rapid estimation of epidemiological parameters is essential to establishing appropriate public health measures [1,3]. Inference methods in epidemiology are under rapid development as a result [4–7].

With the advent of affordable sequencing techniques, infected individuals can now be sampled in order to sequence genes (or even the complete genome) of the pathogen causing their infection. In the case of outbreaks, this sampling can represent a significant proportion of infected hosts. For instance, Gire et al. sequenced the whole genome of viruses from 78 Ebola cases sampled from late May to mid June 2014 and published a complete analysis of the early spread of this large Ebola outbreak by mid-September [8]. It was estimated that these samples represented a significant proportion of all the infections in Sierra Leone that had taken place at that time [9, 10].

A dated phylogeny can readily be inferred from virus sequences with a known sampling date. Such a ‘genealogy’ of infections bears many similarities with a transmission chain and potentially contains information about the spread of the epidemic. This idea was popularised by Grenfell et al. [11], who coined the term ‘phylodynamics’ to describe the hypothesis that the way rapidly evolving parasites spread, leaves marks in their genomes and in the resulting phylogeny.

Obtaining quantitative estimates from phylogenies of sampled epidemics remains a major challenge in the field [12, 13]. In most studies, epidemiological parameters are inferred using a Bayesian framework based on a likelihood function that describes the probability of observing a phylogeny given a demographic model for a set of parameter values. This model is also referred to as the ‘tree prior’ [14]. Epidemiological dynamics were first captured in the tree prior by using coalescent theory and assuming an exponential growth rate of the epidemic [15], or more flexible variations in the effective population size over time (i.e. effective prevalence) [16–18].

More recently, much progress has been made in deriving tree priors more relevant to epidemiological models (see [19] for a review). In 2009, Volz et al. [20] managed to express the likelihood function of SIS (for ‘Susceptible-Infected-Susceptible’) and SIR (for ‘Susceptible-Infected-Removed’) epidemiological models using coalescent theory, thus allowing for the estimation of *R*_0_. One year later, Stadler [21] derived the likelihood function of a phylogeny using the birth-death process. The method was then extended to other epidemiological models and now allows for the inference of both *R*_0_ and the duration of the infection [22, 23]. Other developments have combined state-of-the-art techniques in epidemiological modelling, for instance particle filtering, with the coalescent model for phylodynamics inference [24–26]. The success of these tree priors was made possible by advances in computing power, and the generalisation of computationally intensive techniques to explore the parameter space, such as Markov Chain Monte Carlo (MCMC) procedures [27]. Many of the tree priors and procedures described above, are implemented in the software package BEAST [14].

Since all the aforementionned phylodynamics studies rely on estimating the likelihood function of a phylogeny under a given epidemiological model, they share the same two limitations. First, it is challenging to express the likelihood function for models that are more realistic than the simple SI or SIR models. Second, even when the likelihood function can be expressed, challenging numerical problems remain. For instance, some analytical formulations (especially integrals) are extremely complicated to solve numerically [23]. Furthermore, size matters and current Bayesian methods have already shown their difficulties to compute likelihood functions for medium size phylogenies, i.e. with more than a few hundreds tips [9, 28], while data sets and trees with thousands strains are not uncommon (e.g. [29, 30]).

The likelihood-free method, Approximate Bayesian Computation (ABC), may offer a means to circumvent both these limitations. In short, ABC proposes to bypass the difficulty in computing (and even sometimes formulating) the likelihood function, by performing simulations and comparing the simulated and ‘target’ data [31–34]. This comparison is made possible through the description of the data by summary statistics. In the end, as the name suggests, the likelihood is approximated instead of being computed.

By summarising phylogenies via summary statistics, one can then calculate a distance between the simulated and target (observed) phylogenies. The basic ABC algorithm, called rejection [35], consists in retaining a small fraction of simulations that are close to the target in view of the computed distances. The epidemiological parameters that correspond to these retained simulations, constitute the final posterior distribution of the parameters. Over the last decade, several improvements of the rejection algorithm have been proposed. The first is a reference algorithm aimed at making the search in the parameter prior space more efficient, for instance by using MCMC (ABC-MCMC) [36]. Newer ABC methods further adjust the posterior distribution obtained by rejection. This can be done using Sequential Monte Carlo (ABC-SMC), which consists in re-sampling parameters from the posterior and thus iterating the rejection process until convergence [37, 38]. An alternative consists of adding a regression step (linear or not) [35, 39]. In the latter method, the simulations selected by rejection are used to learn a regression model, which is then used to project the simulation distribution on the target. The regression-based ABC has the advantage of being potentially less computationally intensive and also less sensitive to the curse of dimensionality of the set of summary statistics than the ABC-MCMC or ABC-SMC [39].

ABC has been compared to a Bayesian method based on the exact likelihood implemented in BEAST and has been shown to infer epidemiological parameters from genetic data as accurately as and more effectively than the Bayesian method [40]. Yet, to our knowledge, ABC has only been applied to phylodynamics in two studies [41, 42]. As shown in the first study, this could be due to the fact that the approach can be sensitive to the choice of summary statistics and requires careful calibration of the tolerance parameter [41]. More recently, an ABC-MCMC algorithm using a tree shape kernel distance has been developed for epidemiological parameter estimation assuming the Birth-Death-Susceptible-Infectious-Removed model (BDSIR [28]) and showed promising results [42].

Here, we further assess the accuracy of regression-based ABC approaches to infer epidemiological parameters from phylogenies. To this end, we first designed a large variety of summary statistics (83 in total) on both the phylogeny and its associated lineage-through-time plot. The choice of such a large number of statistics was motivated by the existence of powerful machine learning techniques that perform regression using variable selection, such as the regularized neural network model introduced by Blum [39]. This also allowed us to analyse where the epidemiological information was located in the phylogeny. We considered several epidemiological scenarios in order to have a broad and accurate view of the performance of ABC methods to infer epidemiological parameters. In particular, we used two classical epidemiological models: the Birth-Death (BD) model and the Susceptible-Infectious-Removed (SIR) model. For the former model (BD), the exact likelihood function of the phylogeny is known [21, 22, 43], whereas for the latter model (SIR), it is typically approximated. Indeed, the likelihood function of the SIR model can be derived analytically [23] but its numerical computation for large population size requires simplifying assumptions that lead to the BDSIR model [28]. Lastly, to further assess the power of our method, we re-analysed data from the early stages of the recent Ebola epidemic in Sierra Leone [8–10]. Overall, we found that the accuracy of the estimates obtained using regression-based ABC approaches are comparable to that based on the likelihood function. ABC can even outperform existing methods with regards to accuracy and computing time for more complex models and large phylogenies, making it more valuable as the size of infection phylogenies increases.

## Materials and Methods

### Compartmental models

We considered three epidemiological models: a Birth-Death (BD) model (Fig. 1a), a Susceptible-Infected-Removed (SIR) model without demography (i.e. with a constant host population size, Fig. 1b) and a Birth-Death model with an Exposed class (BDEI, Fig. 1c). The BD model [43] and an approximate version of the SIR model, the BDSIR model [28], have been implemented in BEAST2 [44]. The BDEI model has been used for likelihood-based parameter inference of the early spread of Ebola epidemics in Sierra Leone [9]. These compartmental models are defined by ordinary differential equation (ODE) systems (see Supplementary Text S1).

**Figure 1.**
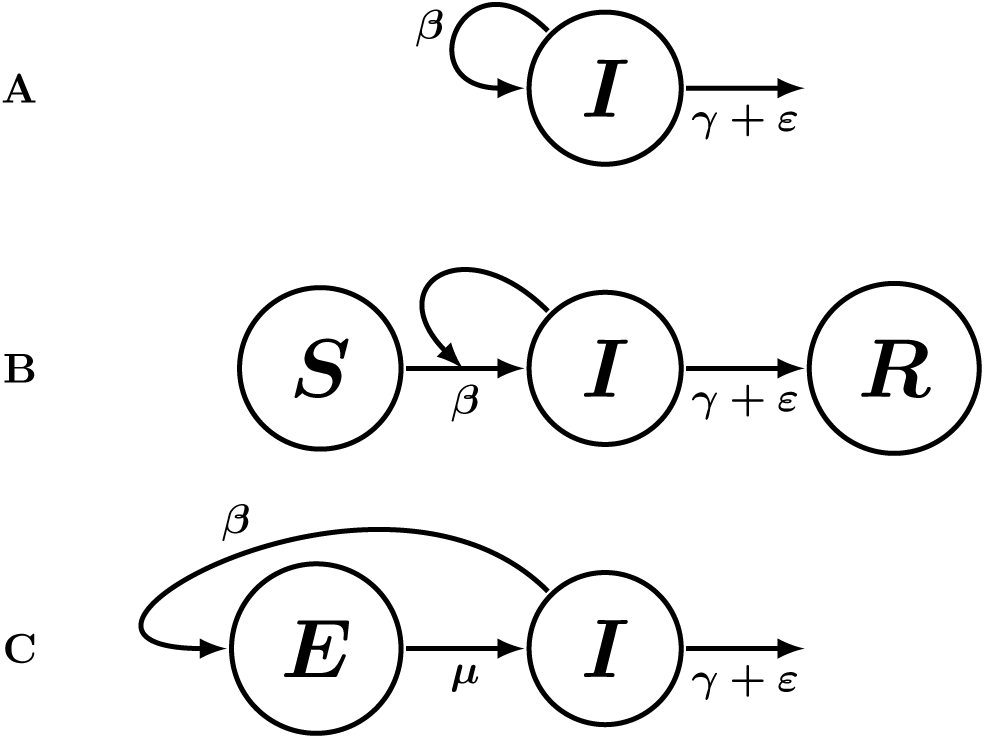
The epidemiological models. (a) The Birth-Death (BD) model. (b) The Susceptible Infected Removed (SIR) model. (c) The Birth-Death model including an Exposed class (BDEI). The four compartments correspond to susceptible (*S*), exposed (*E*), infectious (*I*) and removed (*R*) individuals. In BD and BDEI models, new infections arise at a constant (‘birth’) rate *β* per infectious individual. In the SIR model, the number of new infections depends on the number of susceptible individuals, the transmission rate *β* and the number of infectious individuals. In this latter model, the total host population size is assumed to be constant (*N*). In all models, infections end (i.e. ‘die’) at a rate γ + *ε*.

In these models [2], individuals susceptible to the pathogen become infected after a contact with infectious individuals and a successful transmission, which occurs at an overall transmission rate *β*. Following infection, individuals either become infectious immediately (BD and SIR models) or at a rate *μ* after a latency period in the Exposed class (BDEI model). They are then ‘removed’ (i.e. recover with a lifelong immunity or die) at a rate γ. Finally, they can be sampled, at a rate *ε*. By sampling, we mean that the pathogen is sequenced from the patient. Because sampling generally leads to treatment or at least to behavioural changes, we assumed that infected individuals are also ‘removed’ after sampling. This assumption is commonly made in phylodynamics [22, 28, 43] and we kept it here to facilitate comparisons, but it could easily be relaxed. The sampling proportion *p* is defined as the ratio of the sampling rate (*ε*) over the total removal rate (γ + *ε*).

The critical difference between BD models and the SIR model, lies in the transmission rate per infected individual λ(*t*): this rate is constant in the BD models (λ(*t*) = *β*), but it depends on the susceptible population size in the SIR model 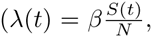 where *S*(*t*) is the number of susceptible individuals at time *t* and *N* is the effective population size). In other words, the SIR model assumes an effective host population with a fixed size *N* and which is initially fully susceptible (*S*(*t* = 0) = *N*). The susceptible population is depleted as the epidemics spreads (*S*(*t* > 0) < *N*), and this depletion decreases the speed of the spread of the epidemics (λ(*t* > 0) < λ(*t* = 0)).

In this study, our goal is to infer, from dated phylogenies, the vector of epidemiological parameters *θ* composed of:

- the mean duration of the infectious period 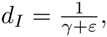
- for the BDEI model, the mean duration of the latency period 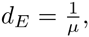
- the basic reproduction number, or the number of secondary cases, which is defined as 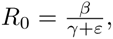
- and, for the SIR model, the effective population size *N* = *S* + *I* + *R*.

Contrary to the likelihood-based phylodynamics methods [9,28,43], we did not attempt to infer the sampling proportion using ABC, since γ and *ε* are inseparable in the epidemiological models we studied in this paper (see Supplementary Text S1) [45].

### Epidemiological Event-Driven Model (EEDM)

The compartmental models described above are deterministic continuous-time models. However, whatever the method used (likelihood-based or not), epidemiological parameter inference requires taking into account the stochasticity of events at the level of the individual in the real-world epidemics. This is done here by implementing an event-driven version of the ODE-based models.

A dated phylogeny of an epidemic can be viewed as a genealogy of infections where each branching represents a transmission and each leaf an end of infection. To simulate such an object, one needs a discrete-time stochastic simulation algorithm. Gillespie’s Direct Algorithm is an event driven approach that is commonly used to simulate chronologies of epidemiological events [2,46], sometimes called trajectories [47], assuming a compartmental model. The translation of a compartmental model into an event-driven model via Gillespie’s algorithm requires the specification of all events that may occur. In BDEI model for instance, these events are: ‘transmission’, ‘end of latency’, ‘removal’ and ‘sampling’, which occur respectively at rates *β*, *μ*, γ, and *ε*, per infected individual. Note that a great advantage of this algorithm is that there is an exact correspondence between the stochastic simulations and the deterministic (continuous and ODE-based) model.

It is possible to build a tree while simulating epidemiological dynamics by using the analogy between a genealogy of infections and a transmission chain [11]. Practically, this means assuming that each transmission event leads to a branching in the tree. In the BDEI model, after transmission, an exposed individual is not infectious yet and the new branch is said to be ‘passive’ because it cannot perform branching. It is only if an ‘end of latency’ event occurs that a passive branch is activated, thereby allowing it to branch. Lastly, when a ‘removal’ or a ‘sampling’ event occurs, a leaf is created because the infection ends.

The simulation of an EEDM produces a full transmission tree. However, pathogen sequences are only available for infected individuals who are ‘removed’ by sampling. To perform comparisons with a dated phylogeny, we need a simulated tree of a sampled epidemics and thus removed all the phylogenetic history of the non-sampled individuals.

### Summary statistics

Dated phylogenies are complex objects. Therefore, to compare them and capture the epidemiological information they may contain, we used summary statistics. We decided to compute as many summary statistics as possible to capture as much information as possible. This was motivated by the fact that there is no consensus in the field regarding which summary statistics to use. Importantly, this decision was made possible by the existence of efficient regression models that perform variable selection and can be combined to ABC (see below). Overall, we used 83 summary statistics which we group into ‘families’ to better identify where the epidemiological information is in the phylogeny.

These involve objects such as branch lengths (Tab. 1), tree topology (Tab. 2) and the Lineage-Through-Time (LTT) plot (Tab. 3) [48].

**Table 1.** Summary statistics based on branch lengths (bl set) • Statistics computed on three time-based parts of the tree. Internal branches belong respectively to the first (*k* = 1), second (*k* = 2) or third (*k* = 3) part of the tree if they end before the first, second or third delimitation, respectively. ^÷^ Ratios between each piecewise statistic related to internal BL and the same statistic computed on all external BL.

| Notation | Description |
| --- | --- |
| $max\_H$ | Sum of the branch lengths between the root and its farthest leaf |
| $min\_H$ | Sum of the branch lengths between the root and its closest leaf |
| $a\_BL\_mean$ | Mean length of all branches |
| $a\_BL\_median$ | Median length of all branches |
| $a\_BL\_var$ | Variance of the lengths of all branches |
| $e\_BL\_mean$ | Mean length of external branches |
| $e\_BL\_median$ | Median length of external branches |
| $e\_BL\_var$ | Variance of the lengths of external branches |
| $i\_BL\_mean\_ [k]^\bullet$ | Piecewise mean length of internal branches |
| $i\_BL\_median\_ [k]^\bullet$ | Piecewise median length of internal branches |
| $i\_BL\_var\_ [k]^\bullet$ | Piecewise variance of the lengths of internal branches |
| $ie\_BL\_mean\_ [k]^\div$ | Ratio of the piecewise mean length of internal branches over the mean length of external branches |
| $ie\_BL\_median\_ [k]^\div$ | Ratio of the piecewise median length of internal branches over the median length of external branches |
| $ie\_BL\_var\_ [k]^\div$ | Ratio of the piecewise variance of the lengths of internal branches over the variance of the lengths of external branches |

**Table 2.**
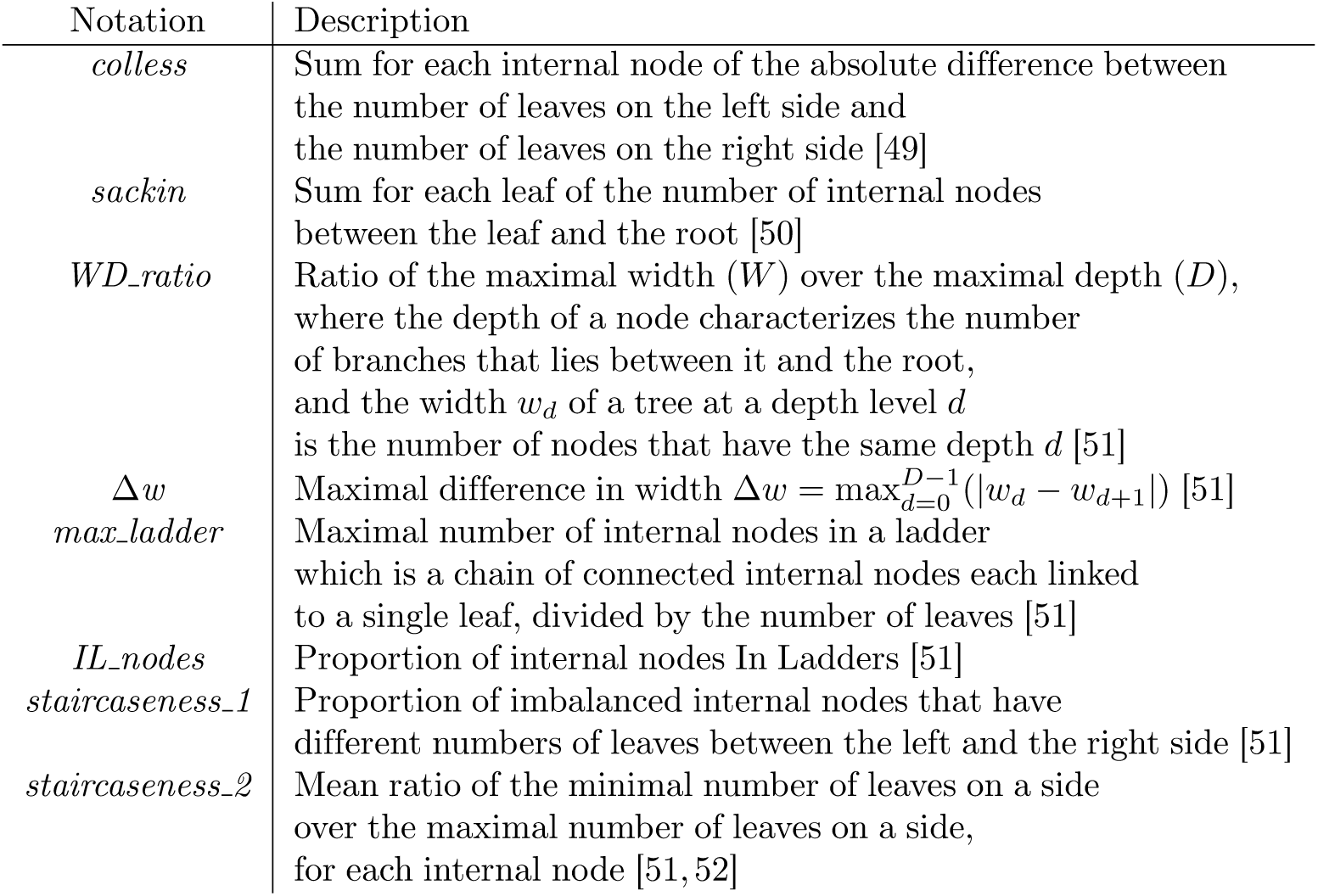
Summary statistics based on the tree topology (topo set).

**Table 3.**
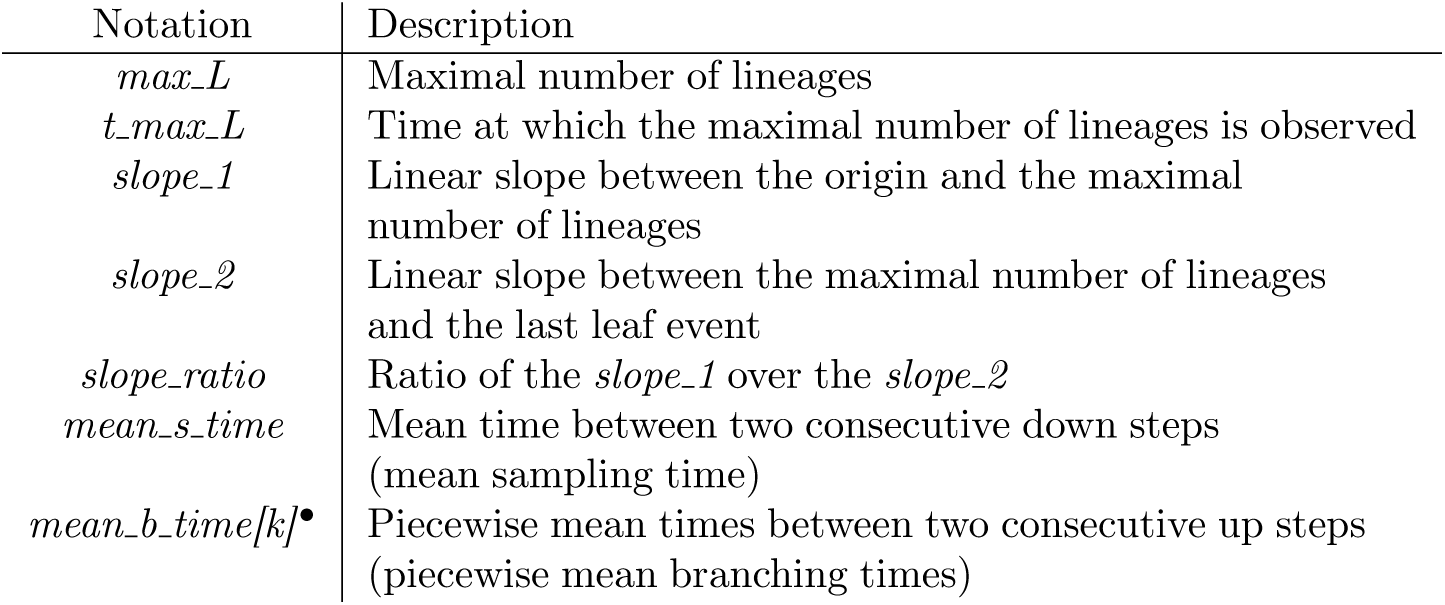
Summary statistics based on the LTT plot (ltt set) •Computed on three part of the tree. Consecutive steps up respectively to the first (*k* = 1), second (*k* = 2) or third (*k* = 3) part of the tree if the second steps happens before the first, second or third delimitation, respectively.

Since branching occurs throughout the phylogeny at a rate that varies through time (the number of infected hosts can vary and also the number of susceptible hosts in the SIR model), we designed all the summary statistics related to branching and internal branches (linking two internal nodes) in a piecewise manner (Tab. 1). We temporally cut the tree into three equal parts: internal branches belong respectively to the first, second or third part of the tree, if they end before the first (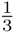max_*H*), second (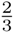*max_H*) or third (*max_H*) delimitation, respectively, where *max_H* represents the height of the farthest leaf.

The sampling rate also varies through time. However, sampling events often appear late in the phylogeny, which is why we only computed global (on the whole tree) summary statistics to describe sampling events and external branches (linking internal nodes to the leaves).

It is known that the topology of a phylogeny can be driven by processes such as immune escape [11]. Moreover, it has been shown recently that different transmission patterns can result in quantitatively different phylogenetic tree topologies. In particular, heterogeneity in host contact can influence the tree balance [51]. This is why we also used phylogenetic topological indexes as summary statistics (Tab. 2).

The Lineage-Through-Time (LTT) plot provides a graphical summary of a phylogeny [48]. It represents the number of lineages along the phylogeny as a piecewise constant function of time (Fig: 2). Each step up in the LTT plot corresponds to a branching in the phylogeny, and each step down to a leaf. If all infected individuals of an epidemics are sampled, the phylogeny corresponds to the full transmission tree and the LTT plot is identical to the prevalence curve. Therefore, as noted in earlier studies [23,53-55], it is reasonable to think that this plot could contain information about the epidemiological parameters. We summarized this plot with two sets of summary statistics, one that captures particular measures of the LTT plot (Tab. 3) and another that simply uses the coordinates of its points as ‘summary’ statistics. For this latter set of summary statistics, because the LTT plot contains as many points as there are nodes in the phylogeny (a phylogeny of *n* leaves has 2*n* - 1 nodes so its LTT plot has 2*n* - 1 points) and because we here consider phylogenies with more than 100 leaves, we averaged the points into 20 equally-sized bins, thus generating 40 summary statistics (20 x-axis coordinates and 20 y-axis coordinates).

**Figure 2.**
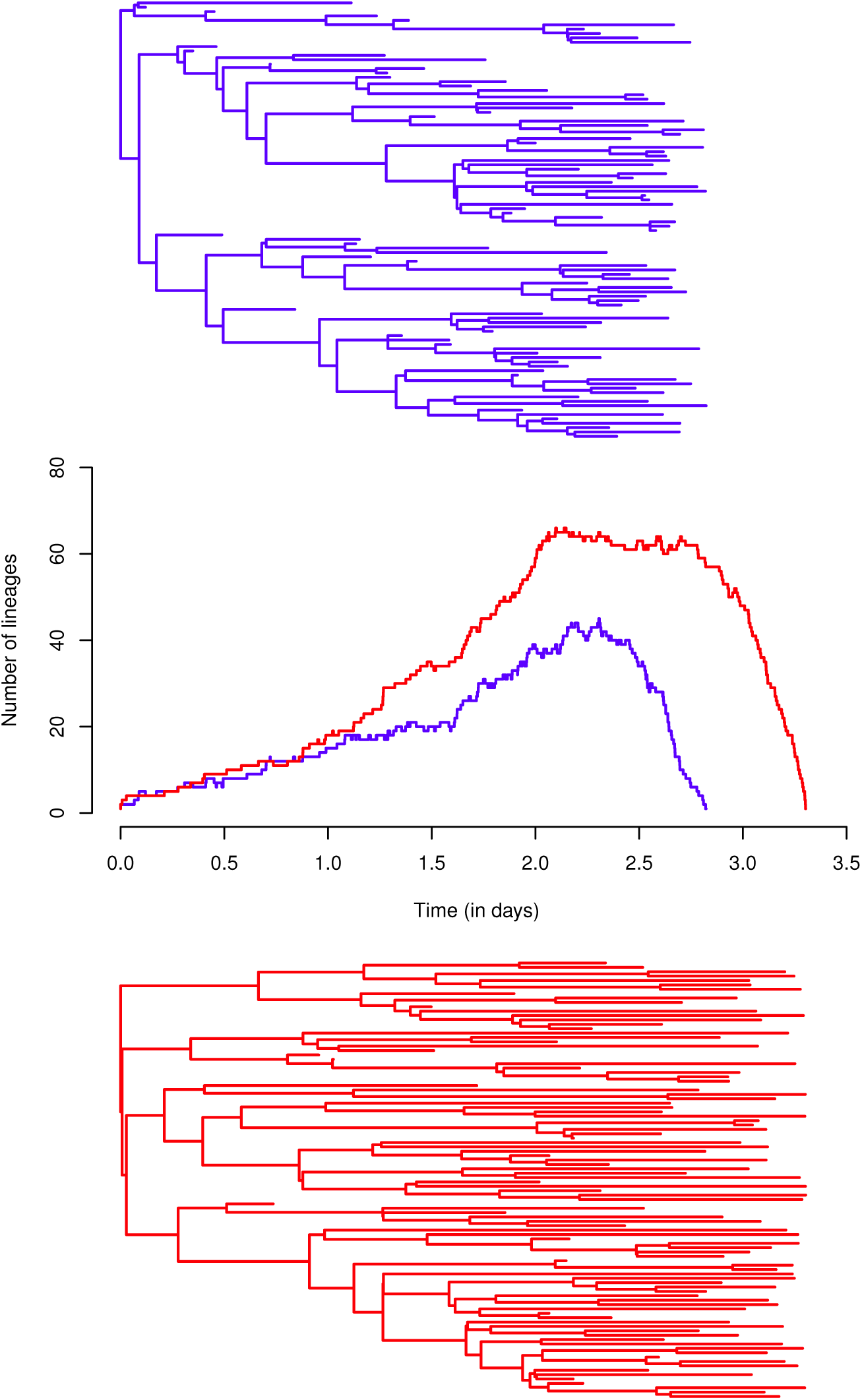
Simulated phylogenies of 100 leaves assuming a BD model and their corresponding LTT plot. The red phylogeny was simulated assuming *θ* = (*R*_0_ = 10, *d*_*I*_ = 5, *p* = 0.5) and the blue phylogeny was simulated assuming *θ* = (*R*_0_ = 2, *d*_*I*_ = 5, *p* = 0.5). Different *R*_0_s lead to different LTT plots and different tree shapes.

To sum up, we used two main sets of summary statistics:

- the sUMsTATs set, with 43 summary statistics related to the tree and its LTT plot
  - the ToPo set: 8 topology summary statistics,
  - the BL set: 26 branch-length summary statistics,
  - the LTT set: 9 summary statistics related to the LTT plot,
- the coords set, with 40 mean coordinates of the LTT plot.

### Simulation study

We wanted to assess the potential of ABC methods to infer epidemiological parameters from phylogenies. To this end, we compared ABC methods and Bayesian methods involving the derivation of the likelihood function of the phylogeny (hereafter referred to as ‘Bayesian methods’) on ‘target’ trees that had been simulated
under a variety of scenarios. In particular, we used the BD and the SIR epidemiological models to perform exhaustive comparisons. We expected our method to perform less well than Bayesian methods since ABC, by definition, only approximates the likelihood using simulations. However, practically, the implementation of likelihood-based approach often requires simplifying assumptions to allow for efficient computation, which makes the results of the comparison non trivial to predict.

**Target trees**. We considered 32 scenarios, which correspond to all the combinations between:

- 2 epidemiological models (BD and SIR),
- 2 *R*_0_ values (*R*_0_ = 2, for a slow Influenza-like spread, and *R*_0_ = 10, for a rapid Measles-like spread),
- 2 durations of infection (*d*_*I*_ = 5 and *d*_*I*_ = 30),
- 2 sampling proportions (*p* = 0.05 and *p* = 0.5),
- 2 tree sizes (100 leaves and 1,000 leaves),

SIR target trees were all simulated in a population with *N* = 25, 000 individuals.

To carry out a statistical performance analysis we simulated 100 phylogenies (replicates) for each scenario.

### Simulated ‘training’ trees for ABC

For each of the 32 sets of simulated target trees, we simulated a set of 10, 000 trees to train the ABC. We assumed the values of all the epidemiological parameters to be distributed in uniform priors (see Tab. 4).

**Table 4.**
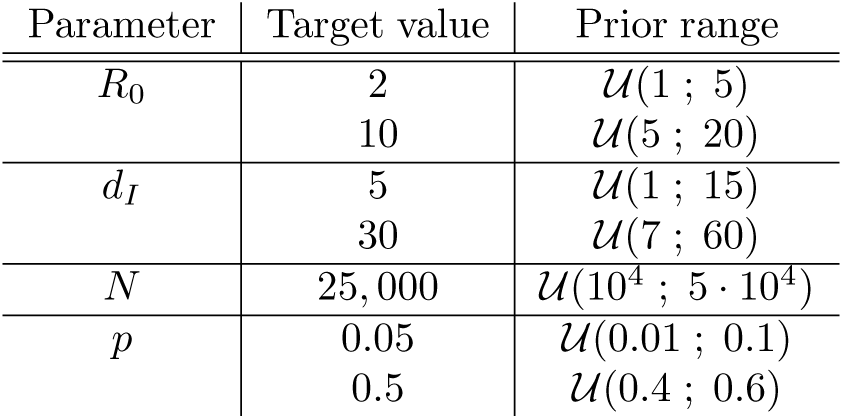
Prior table.

### Correlation analysis

After simulating trees and computing the 83 summary statistics on each tree, we looked for where the epidemiological information was in the trees. We used Spearman’s correlation between each of the summary statistics and epidemiological parameters.

### ABC

We used the *abc* function from the abc R package [39,56] to infer posterior distributions from rejection alone, and rejection followed by adjustment using feed-forward neural network (FFNN). This function performs the rejection algorithm of Beaumont et al. [35] using a tolerance parameter *P*_*δ*_, which represents a percentile of the simulations that are close to the target. The proximity of the simulations to the target is evaluated in the function via the euclidean distance between each normalized simulated vector of summary statistics, and the normalized target vector. The acceptance region is therefore spherical.

Prior to adjustment, the *abc* function performs a smooth weighting using an Epanechnikov kernel as for the loc-linear adjustment proposed by Beaumont et al. [35]. We then performed an FFNN adjustment using the option available in the *abc* function [56]. This adjustment involves the construction of a non-linear conditional heteroscedastic regression model, using the *nnet* function (nnet R package), which involves a FFNN with a single-hidden-layer [39]. The *nnet* function includes a regularization of the fitting criterion through a penalty on the ‘roughness’. This penalty, called weight decay, corresponds to the sum of the squares of the weights put on the links of the neural network. This penalty contributes to avoid over-fitting [57]. Bishop [58] also states that choosing a number of hidden units lower than the number of variables leads to dimensionality reduction and smoother regression. We used the default parametrization of the *abc* function, which does not provide a perfect control over the regularization, and uses 5 FFNN hidden units.

In addition to simple rejections (ABC) and to rejections with non-linear adjustment using FFNN (ABC-FFNN), we also used a linear adjustment with variable selection using Least Absolute Shrinkage and Selection Operator (LASSO) regression [59]. Using such a regression model that performs a well-controlled dimensionality reduction was motivated by the high number of summary statistics.

We implemented the LASSO adjustment (ABC-LASSO) using the glmnet R package [60]. As in the ABC-FFNN method, we weighted the simulations retained by rejection using an Epanechnikov kernel and we corrected for heteroscedasticity. The variable selection in LASSO is based on the selection of the first components of a principal component analysis [59]. The number of components selected was adjusted using cross-validation with the *cv.glmnet* function. A multi-response gaussian LASSO model was then computed using the *glmnet* function. The information about the variable selection was kept to see whether some specific summary statistics are more often selected than others.

For completeness, we also performed a rejection that uses a functional distance (ABC-D): the distance between two LTT plots. This use of the distance between two non-normalized LTT plots was inspired by the function *nLTTstat* (nLTT R package), which computes the difference between two normalized LTT plots [61]. However, we did not normalize the LTT plots, to account for the potential temporal shift between two LTT plots.

We ran all ABC methods (ABC, ABC-D, ABC-FFNN and ABC-LASSO) to estimate the parameters of all target trees, using the sumstats and coords sets of summary statistics together or separately. We also used different tolerance proportions *P*_*δ*_ = {0.01; 0.05; 0.1; 0.2; 0.3; 0.4; 0.5} to determine the optimal value for each method.

### Likelihood-based Bayesian inference

We inferred the posterior distributions of the epidemiological parameters of the target trees using the likelihood-based Bayesian approaches implemented in BEAST2 [44]. These methods are often used to infer the phylogeny and the epidemiological parameters from dated DNA sequences simultaneously, but they also allow the user to assume that the phylogeny is known. In order to obtain comparable results, we ran BEAST2 with the same simulated dated phylogenies we used for ABC (see [40] for a similar methodology). We also used the same priors in BEAST2 and in our simulations to train ABC methods. The BEAST2 Markov chains were run for 10^6^ steps for all BD scenarios excepted the four scenarios with large trees and low sampling (1,000 leaves and *p* = 0.05), which required 5 · 10^6^ steps for convergence. For SIR scenarios, we ran chains of 10^7^ steps with 100-leaves trees and chains of 5 · 10^7^ steps with 1,000-leaves trees. For all BEAST2 posterior distributions (BEAST2-BD and BEAST2-BDSIR), we discarded the first 10% of the estimates as a burn-in, and controlled for convergence using the Effective Sample Size measure (ESS) for the epidemiological parameters. We checked that ESS was greater than 200 for *R*_0_ and *d*_*I*_, and greater than 100 for *N* estimated on small trees (see Supplementary Table S1).

### Performance analysis

We measured the median 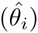 and the 95% Highest Posterior Density (HPD_95%_) boundaries of each parameter posterior distribution (*D*_*i*_). For each ABC run and each simulated scenario (100 target trees), we computed the mean relative error (MRE) as

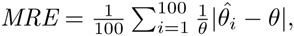

the mean relative bias (MRB) as

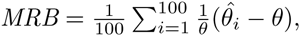

the mean relative 95% HPD width as

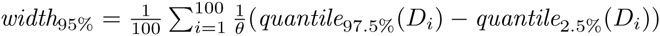

and the 95% HPD accuracy as

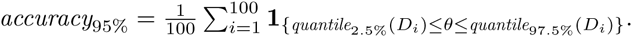

We first tested the influence of the tolerance parameter on the mean relative error (MRE) of the four ABC algorithms (ABC, ABC-D, ABC-FFNN and ABC-LASSO). Then, we compared the performance of all these methods to that of likelihood-based methods implemented in BEAST2, assuming the same models and priors. Lastly, we tested the influence of the epidemiological parameter values used in each scenario on the estimation error (MRE).

### Data analysis: Early stages of the 2014-2015 Ebola epidemic in Sierra Leone

We used the RaxML phylogeny inferred by Gire et al. [8], which was computed on 81 Ebola full-genome sequences: 3 from Guinea patients and 78 from Sierra-Leone patients. The phylodynamics of the Sierra-Leone side of the epidemics has already been investigated by Stadler et al. [9]. To compare our estimates with theirs, we followed their protocol by pruning 6 leaves of the phylogeny corresponding to a sub-epidemics in Sierra-Leone. The remaining 72 sequences were sampled from late May to mid June 2014. Using the known sampling dates, we dated the phylogeny using the Least-Squares Dating (LSD) software, which uses fast algorithms and reaches an accuracy comparable to more sophisticated methods [62].

Stadler et al. inferred epidemiological parameters from this dataset either using BEAST2 with dated sequences (BEAST2-BDEI).

For this data analysis, we assumed a BDEI model and therefore estimated *R*_0_, *d*_*I*_ and the mean duration of latency *d*_*E*_, as in Stadler et al. [9]. As for previous models, the sampling proportion could not be estimated together with the other parameters due to identifiability problems [45].

The Ebola epidemic in Sierra Leone is thought to have started 6 months before it was officially identified and the first sample was collected [8,9]. We therefore needed to consider an additional simulation parameter, *origin*, which is the time before sampling started. Over this time period, the sampling rate was assumed to be *ε* = 0.

We simulated a set of 10,000 ‘training’ trees assuming a BDEI model. For comparison purpose, we first used priors identical to those used in Stadler et al. for their BEAST2-BDEI inferences (see column *p* ≈ 0.7 of Tab. 5). We then used a different interval for the prior on the sampling proportion (*p* ≈ 0.4), because another study suggested that the sampling proportion lies between 0.2 and 0.7 [10]. Moreover, to only simulate biologically realistic epidemiological scenarios [63], we discarded all simulations where the total number of cases went above 50, 000 individuals.

**Table 5.** Prior table for Ebola data.

|  | Assumption |  |
| --- | --- | --- |
| | $p \approx 0.7$ [9] | $p \approx 0.4$ |
| *origin* | $\mathcal{U}(0, 92)$ | |
| $R_0$ | $\mathcal{LN}(0, 1.25)$ | |
| $d_E$ | $\Gamma(0.5, 6)^{-1} \in [1; 26]$ | |
| $d_I$ | $\Gamma(0.5, 6)^{-1} \in [1; 26]$ | |
| $p$ | $\mathcal{B}(70, 30)$ | $\mathcal{B}(25, 35)$ |

As in the simulation study, we computed Spearman’s correlation coefficients between each parameter of the set of simulated trees and each summary statistics.

Rejection is a determinant step in ABC with adjustment because it selects the simulated data that will be used for learning. Even if the chosen regression model is robust, it can collapse if the rejection step fails at retaining a relevant training set. The goodness-of-fit test implemented in the *gfit* function of the abc R package [56,64] is an important preliminary test to be made in data analysis because it indicates whether the summary statistics are informative about the target parameters. This test uses rejection based on the euclidean distance on normalized entries, as defined by Beaumont et al. [35].

Since the dating of the Ebola phylogeny seemed poorly estimated (see Supplementary Figure S1), we performed an upstream test of summary statistics goodness-of-fit of the ‘training’ set against the phylogeny.

We inferred the posterior distributions of *d*_*E*_, *d*_*I*_ and *R*_0_ for the Ebola phylogeny using our ABC-LASSO regression model with *P*_*δ*_ = 0.5. We then compared our own estimates for the epidemiological parameters of the early spread of Ebola epidemic in Sierra Leone with those obtained using the likelihood-based methods of Stadler et al [9]. Finally, we analysed which variables were selected by the LASSO.

## Results

### Locating the epidemiological information in the phylogeny

Figure 3 shows that the 9 summary statistics computed on the Lineage-Through-Time (LTT) plot (LTT set) are the most correlated to the epidemiological parameters of the SIR model. The summary statistics describing the branch lengths (BL set) are less correlated and the topological summary statistics (TOPO set) are, in general, poorly correlated to the parameters. However, the TOPO set becomes more informative when the tree size increases, most likely because topological patterns become more distinguishable. There is little difference in the summary statistics histograms for trees of 100 leaves and trees of 1,000 leaves, the latter being more heavy tailed. BL set summary statistics are positively correlated to the duration of infection (dI) and negatively correlated to the *R*_0_ (see Supplementary Tables S2-S5). None of the topological summary statistics are correlated to *d*_*I*_, even though they are correlated with *R*_0_. The coordinates of the LTT plot that are the most correlated to the epidemiological parameters are those of the x-axis, which are positively correlated to *d*_*I*_ and negatively to *R*_0_. Y-axis coordinates of the LTT plot strongly positively correlate with the *R*_0_ and weakly with the effective population size *N*.

**Figure 3.**
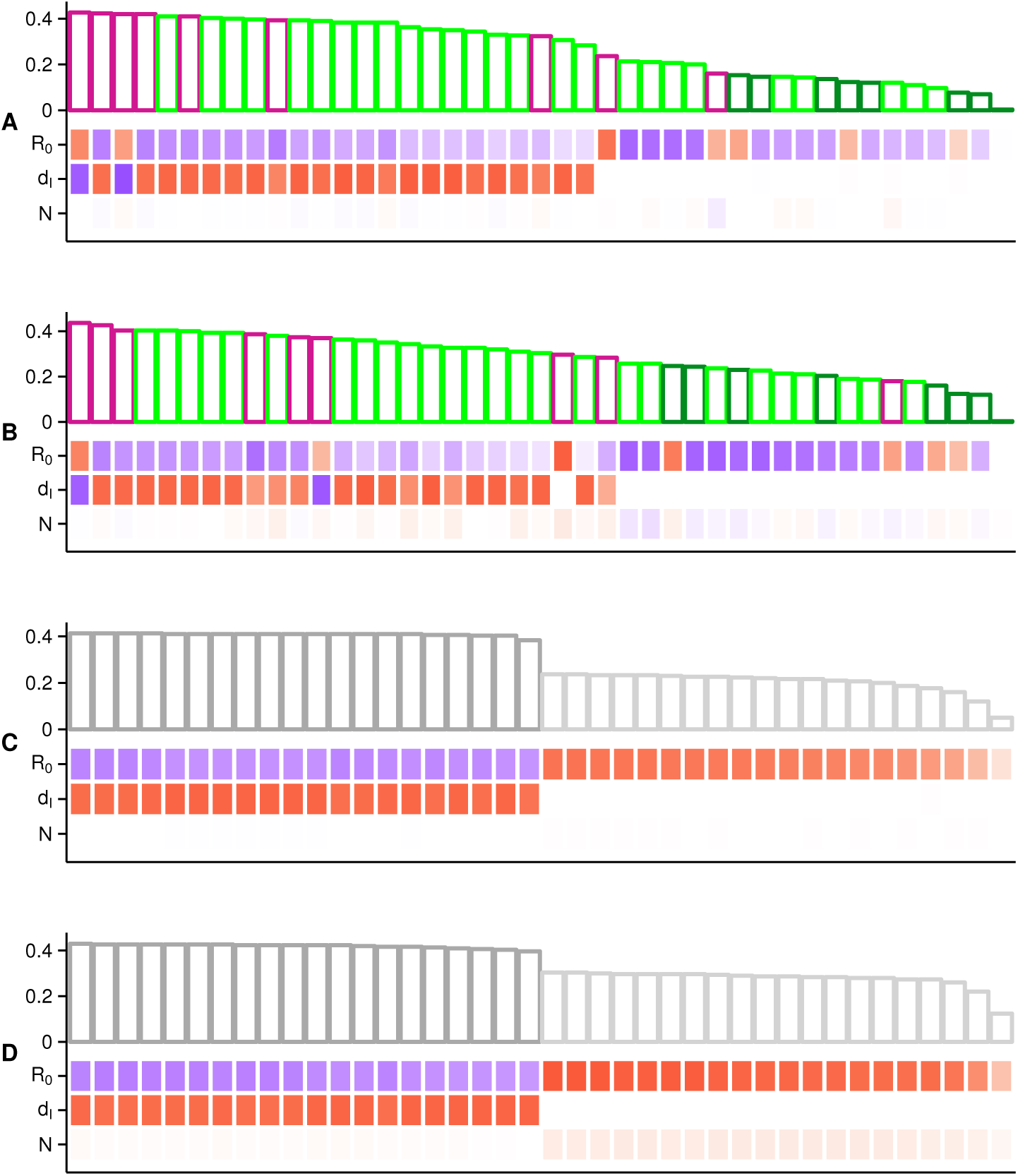
Heat map and histogram of Spearman’s correlations between the epidemiological parameters of the SIR model and all sets of summary statistics for trees of 100 (A and C) or 1,000 (B and D) leaves. In panels A and B, the colors correspond to the bl (light green), topo (dark green) and ltt (magenta) sets. Panels C and D show the coords set related to the LTT plot with x-axis (dark gray) and y-axis (light gray) coordinates. Bar heights in the histograms represent the mean absolute correlation of each summary statistics to the whole set of parameters. Summary statistics and coordinates are ranked from the most to the least correlated. Correlation values between each summary statistics (or coordinate) and each epidemiological parameter are displayed in the heat map, where squares are colored according to a gradient from red (highly positively correlated) to white (no correlation) and blue (highly negatively correlated).

Overall, *R*_0_ is the epidemiological parameter that is the most correlated to all the summary statistics, which suggests that ABC approaches should be able to infer this parameter. On the opposite, Figure 3 bears doubts on the ability of ABC approaches to infer the effective population size from phylogenies, because this parameter is poorly correlated to all summary statistics.

Results for the BD model are very similar to that of the SIR model, except that the mean absolute correlation per summary statistics is increased by 0.2 because of the absence of the *N* parameter in this model (see Supplementary Figure S2 and Supplementary Tables S6-S9).

### Estimating the appropriate tolerance value

In this sub-section, we study the influence of the tolerance parameter used in the rejection step, on the inference error of our four ABC methods: standard rejection (ABC), rejection using the function distance between two LTT plots (ABC-D), rejection and adjustment using regularized neural networks (ABC-FFNN), and rejection and adjustment using LASSO (ABC-LASSO).

We expected the errors of inference of ABC and ABC-D to increase with the tolerance. Indeed, higher tolerance values should cause the rejection step to retain trees that are more and more dissimilar to the target tree, that is, which have been generated by parameter values which are more and more away from the target values. This is what we observe in Figure 4 for *R*_0_ and N when we consider large trees. For the other parameters and for small trees, the errors are similar to the error using the prior (the horizontal gray line), suggesting that there is not enough signal in the summary statistics to infer *d*_*I*_ by ABC and ABC-D.

**Figure 4.**
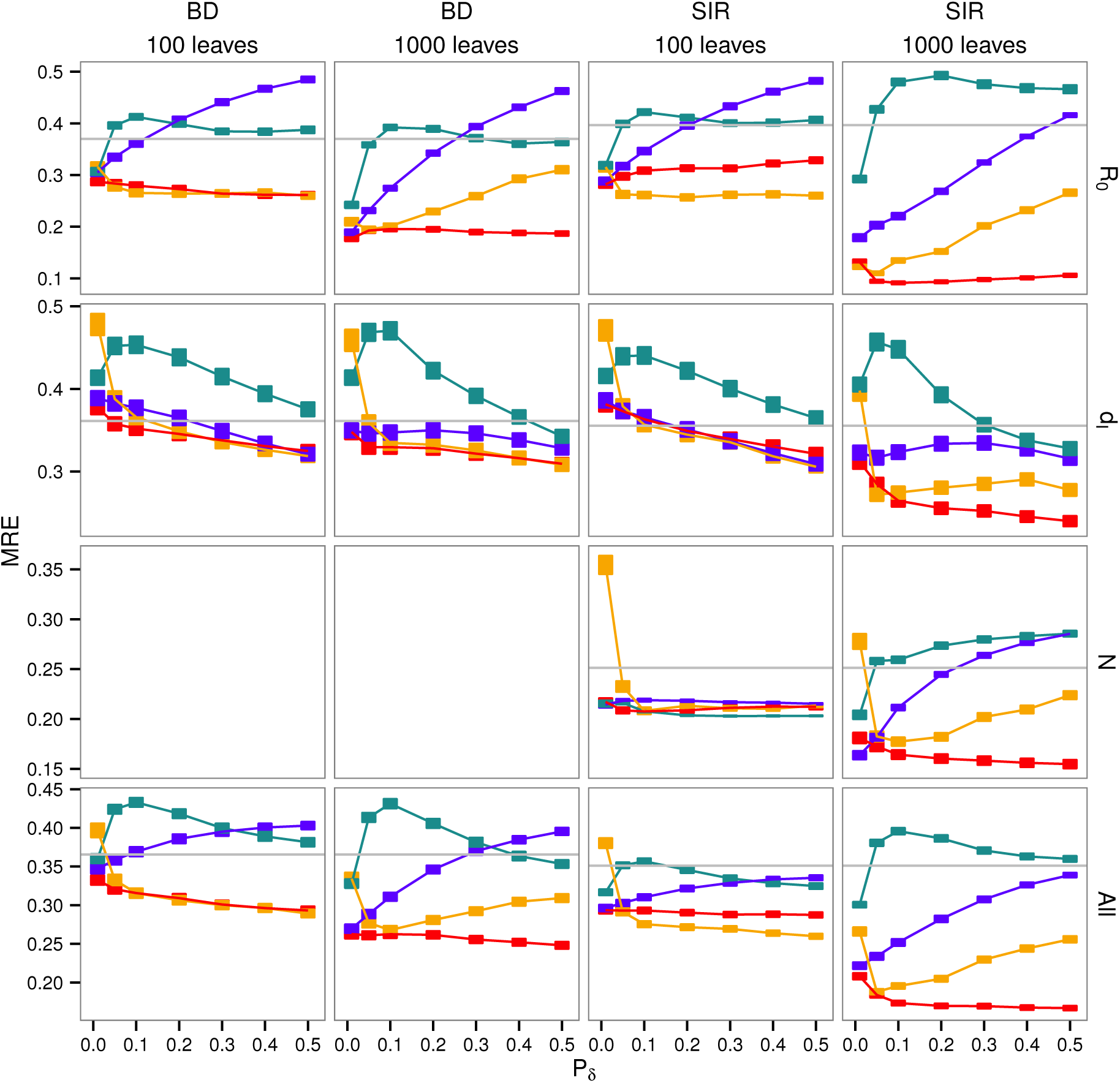
Influence of the tolerance parameter on the error for four ABC approaches used on all summary statistics. The x-axis shows the tolerance value. Squares represent the mean relative errors for each tolerance value with their standard errors. We show errors generated by ABC-D in turquoise, by ABC in blue, by ABC-FFNN in orange and by ABC-LASSO in red. The gray horizontal lines correspond to the mean relative error of the prior (i.e. expected error in rejection with a tolerance of 1). Results are displayed for both BD and SIR model and both trees of 100 leaves and 1,000 leaves.

Regarding the ABC-FFNN method, when the tolerance value increases, we expected the error to decrease at first (because the adjustment method used here requires a certain amount of training data) and finally to reach a plateau (when we have enough data and regularization can control for overfitting effects). This is the case for the inference of *R*_0_ and *N* on small trees and for *d*_*I*_. The error sometimes increases at the end for high tolerance values, which could be due to a poorly controlled regularization or to the limited size of the neural-network in the abc R function.

Concerning the ABC-LASSO method, we expected an increase in the tolerance value to decrease inference error at first for the same reason as for the FFNN. Then, we expected the error to reach a plateau and finally to increase because increasing the size of the training data increases the probability of non-linearity, which is problematic for the LASSO (linear) regression model. This is what we observe in Fig. 4. Furthermore, we see that the relative errors with this method are generally below the threshold represented by the error induced by the prior.

We also analysed the influence of the tolerance parameter on the 95% Highest Posterior Density (HPD) width (*width*_95%_). As expected, the posterior distributions obtained using ABC methods with adjustments are more adjusted than those obtained using the ABC-D or the standard ABC method (see Supplementary Figure S3). The *width*_95%_ of the posteriors obtained using ABC, ABC-D or ABC-FFNN increases with the tolerance. However, the *width*_95%_ of the ABC-LASSO posteriors seems to be insensitive to the tolerance parameter.

Results for the sumstats and coords sets of summary statistics separately are available in Supplementary Figures S4 and S5.

Overall, 0.01 is the best tolerance value for rejections without adjustment and 0.5 is the best value with adjustment. Since this result was observed for both the BD and the SIR model, we adopted these values as default for the rest of the study.

### Performance analysis

Figure 5 shows that for the SIR model, ABC methods often outperform the likelihood-based approach (BEAST2-BDSIR, in black). The inference error (MRE) of the ABC-LASSO (in red) using all the summary statistics, is always below that of the BEAST2-BDSIR with large trees, excepted for *R*_0_ estimation. This can be explained by the fact that the BEAST2-BDSIR assumes an approximation of the true SIR model that speed up MCMC computations. Moreover, in the BDSIR model, the approximation of the number of susceptible individuals through time, *S*(*t*), potentially makes the effective population size *N* hard to estimate [28]. Here, we see that the likelihood-based method largely fails to infer the population size *N* from a uniform prior.

**Figure 5.**
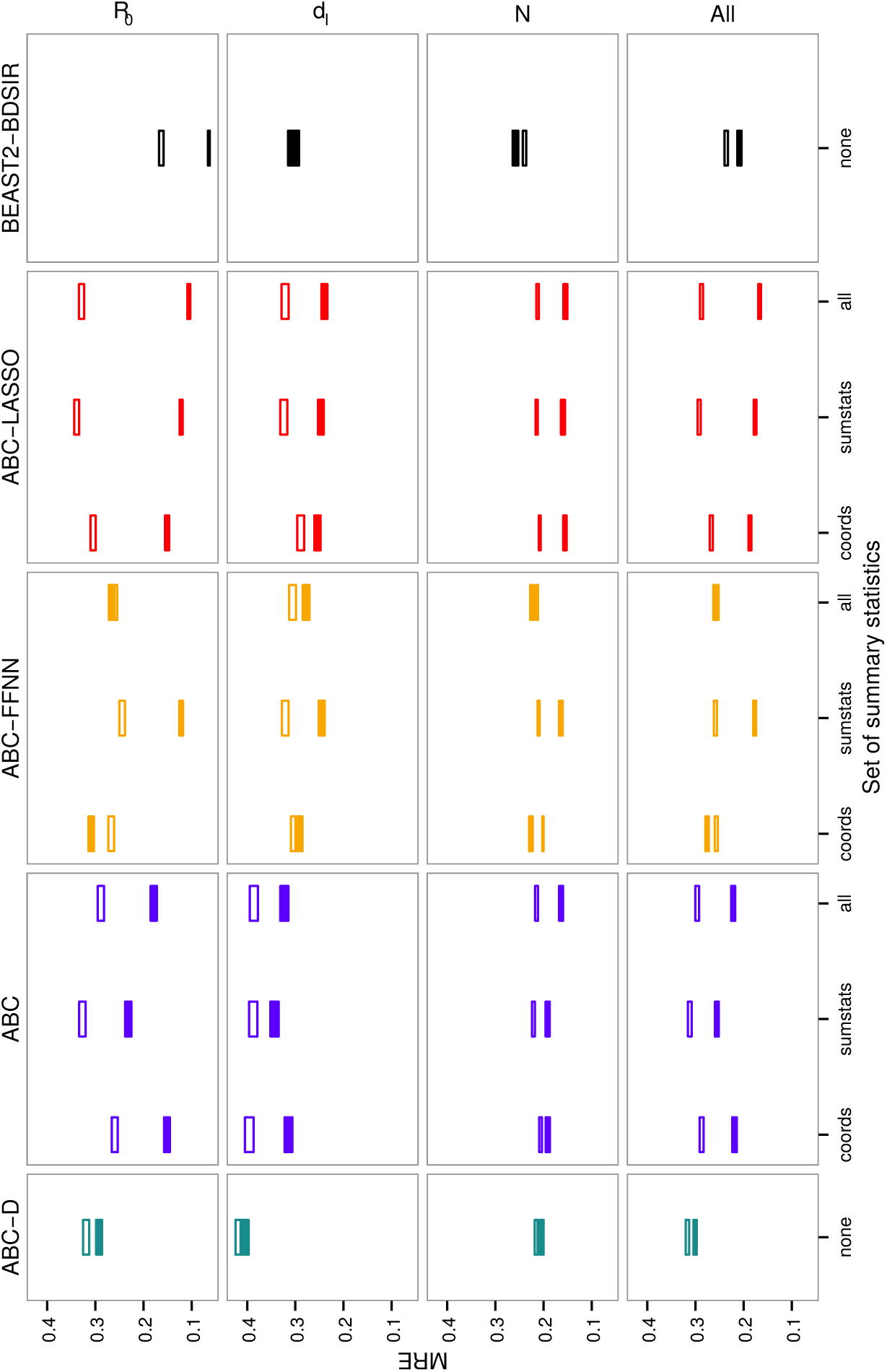
Inference errors on epidemiological parameters of the SIR model using ABC approaches with different sets of summary statistics. The x-axis shows the sets of summary statistics used. Squares represent mean errors with their standard errors. Empty squares correspond to results obtained on trees of 100 leaves and filled squares to results on trees of 1,000 leaves. We show errors generated by ABC-D in turquoise, by ABC in blue, by ABC-FFNN in orange, by ABC-LASSO in red and by BEAST2-BDSIR in black. We show the average errors (bottom row) and the error for each parameter of interest.

The standard ABC method (in blue) already provides good estimations of the *R*_0_. This is consistent the Spearman’s correlations (Fig. 3): the *R*_0_ is the parameter the most correlated to the summary statistics, especially to the coordinates of the LTT plot (coords set).

The standard ABC method (in blue) using the euclidean distance between LTT plot coordinates (coords set), is more accurate than the ABC-D method using the functional distance between two LTT plots (in turquoise). This can be explained by the fact that, in the functional distance, we only consider the differences on the y-axis of the LTT plots, while in the standard ABC using the coords set we also consider the differences on the x-axis, which represents the time variable.

The accuracy of epidemiological parameter inference by ABC-LASSO (in red) is better for all parameters with dated large phylogenies. This is not always the case for the BEAST2-BDSIR method.

The performances of all ABC methods are comparable when we consider small trees. For large trees, ABC-FFNN is the sole method to provide highly variable results, which suggests that the regularization is poorly controlled in the algorithm we used.

ABC-LASSO always gives better estimations than the standard ABC on large trees. It also gives reliable results whatever the set of summary statistics used. This suggests that our LASSO implementation is robust to the high number of explanatory variables.

We analysed which variables were selected in the LASSO regression models but we did not identify any strong selection pattern. This might be explained by the fact that many variables are highly correlated (results not shown). Since the importance of summary statistics from one set to another changes as a function of the epidemiological scenario considered and since our ABC-LASSO is robust to large numbers of variables, we recommend to infer epidemiological parameters using ABC-LASSO with all summary statistics.

Results concerning the BD model are presented in Supplementary Figure S6 and are globally similar to what is observed for the SIR model, except that none of the ABC methods outperform BEAST2-BD. This is consistent with the fact that BEAST2-BD is based on the exact likelihood function of the BD model. Nevertheless, the accuracy of ABC-LASSO on large trees is very close to that of BEAST2-BD.

Inference error is a first way to assess the quality of a fit, but looking at the detailed posterior distribution is more informative. To illustrate this, Figure 6 gives the example of a particular SIR scenario (dense sampling, high *R*_0_ and high *d*_*I*_). For small dated phylogenies (Fig. 6A), we see that posterior distributions of *R*_0_ and *d*_*i*_ obtained from both ABC-FFNN and ABC-LASSO approaches and using all summary statistics are large and similar to the prior distribution. We observe the same pattern for BEAST2-BDSIR, despite the fact that the median of all replicates approximate the target value well. This suggests that there is not enough signal in small trees for reliable estimation. For large dated phylogenies (Fig. 6B), the majority of the replicates of ABC-LASSO converge towards a posterior distribution, which is adjusted and approximatively centered on the target value. This is also true for the BD model (see Supplementary Figure S7). We find similar posterior distributions for the likelihood-based approach except for the *N* parameter, where the posterior clearly reveals a lack of convergence.

**Figure 6.**
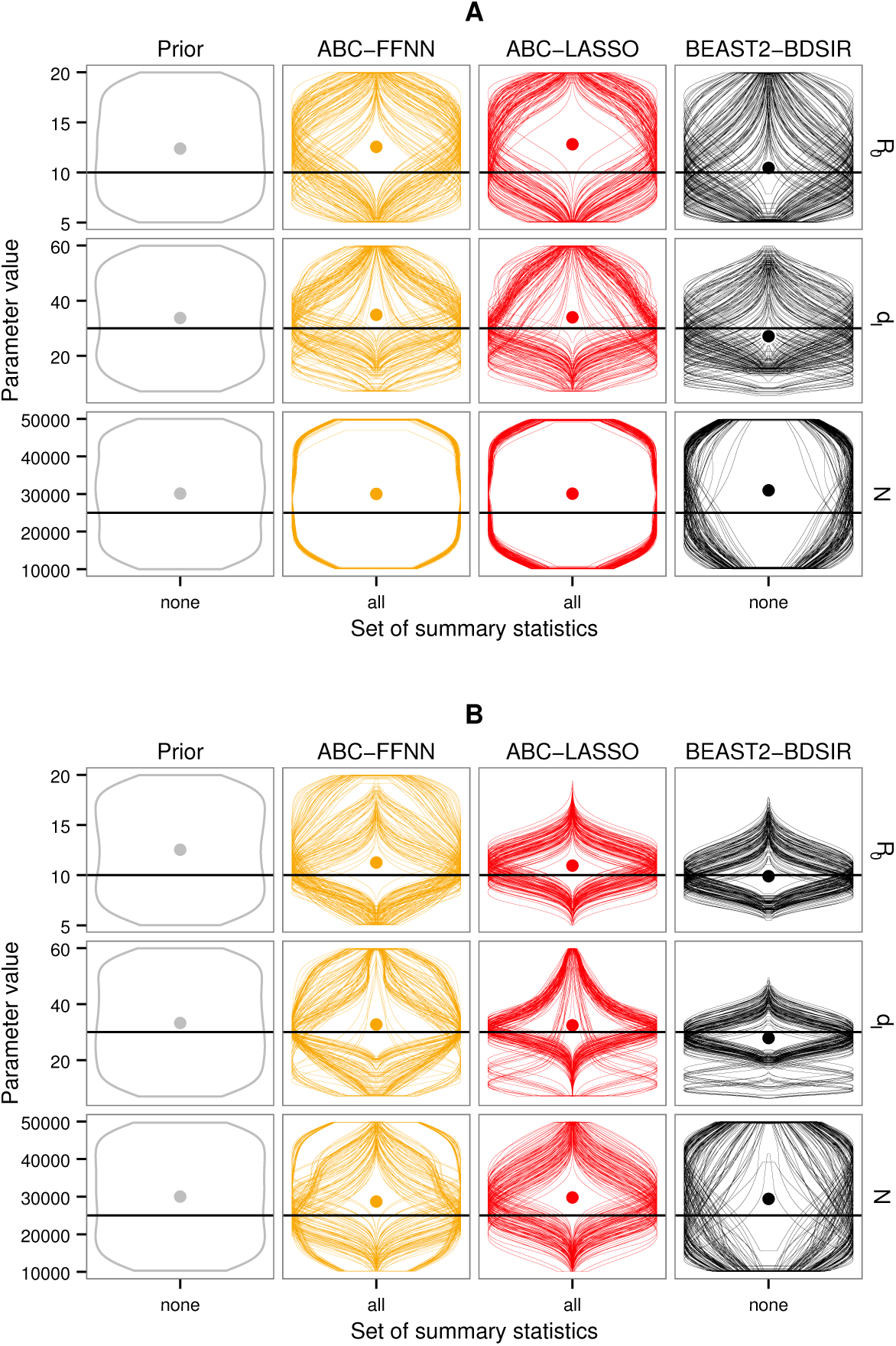
Prior and posterior distributions for parameter estimations by ABC-FFNN, ABC-LASSO and BEAST2-BDSIR. Gray violins correspond to the prior distributions, orange violins to the posterior distributions obtained by ABC-FFNN, red violins by ABC-LASSO and the black violins by BEAST2-BDSIR. All summary statistics were used for both ABC approaches. We displayed the results for one particular epidemiological scenario (*R*_0_ = 10, *d*_*I*_ = 30 and *p* = 0.5) and two different tree sizes: 100 leaves in A and 1,000 leaves in B. There are 100 replicates per panel. The dots represent the median of the merging of the posterior distributions for all replicates. The horizontal black line represents the true value for each epidemiological parameter.

The major advantage of ABC approaches compared to BEAST2 models is that the computation time does not necessary increase with model complexity. In fact, the computation time in our ABC mainly depends on the simulations. For instance, in our simulation study, the ABC-LASSO method ran faster when considering small trees assuming a high sampling proportion (approximately 10 minutes for the BD model and 15 minutes for the SIR model) than when considering large trees assuming a low sampling proportion (approximately 4 hours for the BD model and 2 hours for the SIR model). This time can be considerably reduce if the simulations are ran in parallel. In comparison, BEAST2-BD ran much faster (approximately 1 minute for small trees and, for large trees, 8 minutes assuming high sampling proportion and 38 minutes assuming low sampling proportion). Conversely, BEAST2-BDSIR ran much more slowly (at least 6 hours for small trees and 10 hours for large trees), thus illustrating the limits of the likelihood-based methods for the more complex models. Note that all methods (ABC and BEAST2) estimated the epidemiological parameters assuming a fixed phylogeny.

### Influence of the scenario on inference accuracy

We then tested whether factors such as tree size, epidemiological model and parameter values of the target data, affect the accuracy of the epidemiological parameter inference.

Fig. 7 shows that, as expected, ABC-LASSO infers epidemiological parameters of the SIR model more easily from large phylogenies (1,000 leaves) than from small phylogenies (100 leaves). Surprisingly, the method also performs better for long durations of infection. We expected the sampling proportion (*p*) to have an effect because we thought that a more dense sampling would provide more precise knowledge of the underlying dynamics of the epidemic. This effect was not observed, possibly because of the tight prior on *p*.

**Figure 7.**
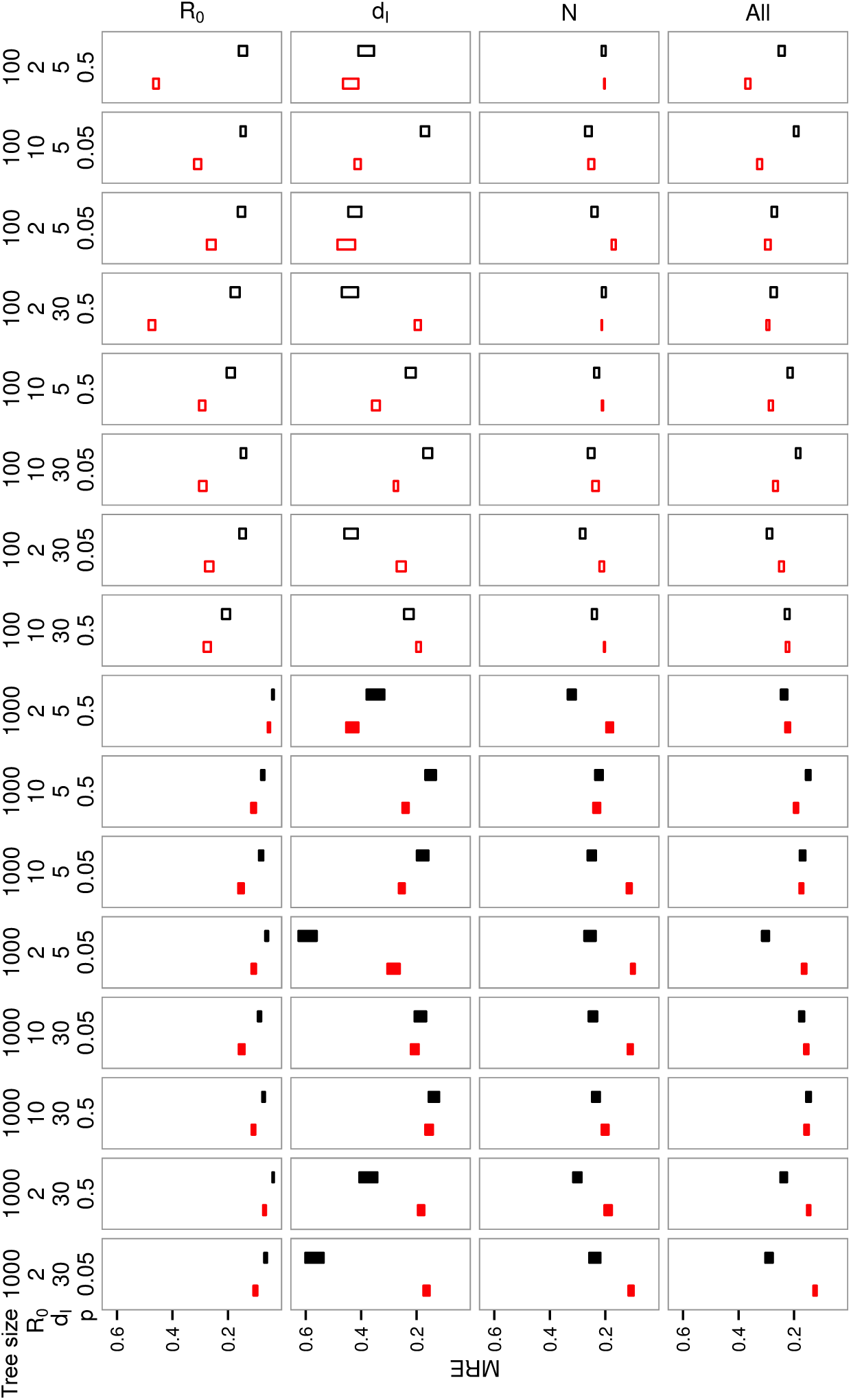
Ranking SIR scenarios based on the error of inference. On the x-axis, scenarios are ordered ranging from the easiest to infer epidemiological parameters using ABC-LASSO, to the worst. Squares represent the mean errors with their standard errors. Empty squares correspond to results obtained on trees of 100 leaves and filled squares to results on trees of 1,000 leaves. The errors generated by ABC-LASSO are in red and that generated by BEAST2-BDSIR are in black.

As mentioned before, the likelihood-based approach does not necessarily provides better estimations with large trees, excepted for the *R*_0_. In fact, the worst scenarios for the likelihood-based method consist in large trees simulated assuming *R*_0_ = 2 and *p* = 0.05.

Results concerning the BD model are available in Supplementary Figure S8.

### Ebola phylodynamics

Fig. 8 shows the correlation results for trees of 72 leaves from the recent Ebola outbreak in Sierra Leone assuming a BDEI model and *p* ≈ 0.4 (see Supplementary Figure S9 for *p* ≈ 0.7). As previously observed for the SIR model, we see that the summary statistics computed on the Lineage-Through-Time (LTT) plot (ltt set) and those computed on the branch lengths (bl set) are the most correlated to the epidemiological parameters of the BDEI model. Conversely, the topological indexes (topo set) contain very few information about the parameters. The bl set summary statistics are positively correlated to both the duration of infectiousness *dI* and the duration of latency *d*_*E*_, excepted the *ie_BL_median_[1]* statistics, which is negatively correlated to *d*_*E*_ while positively correlated to *d*_*I*_ (see Supplementary Tables S10 and S11). The coordinates of the LTT plot (coords set) are poorly correlated to *d*_*E*_ (see also Supplementary Tables S12 and S13).

**Figure 8.**
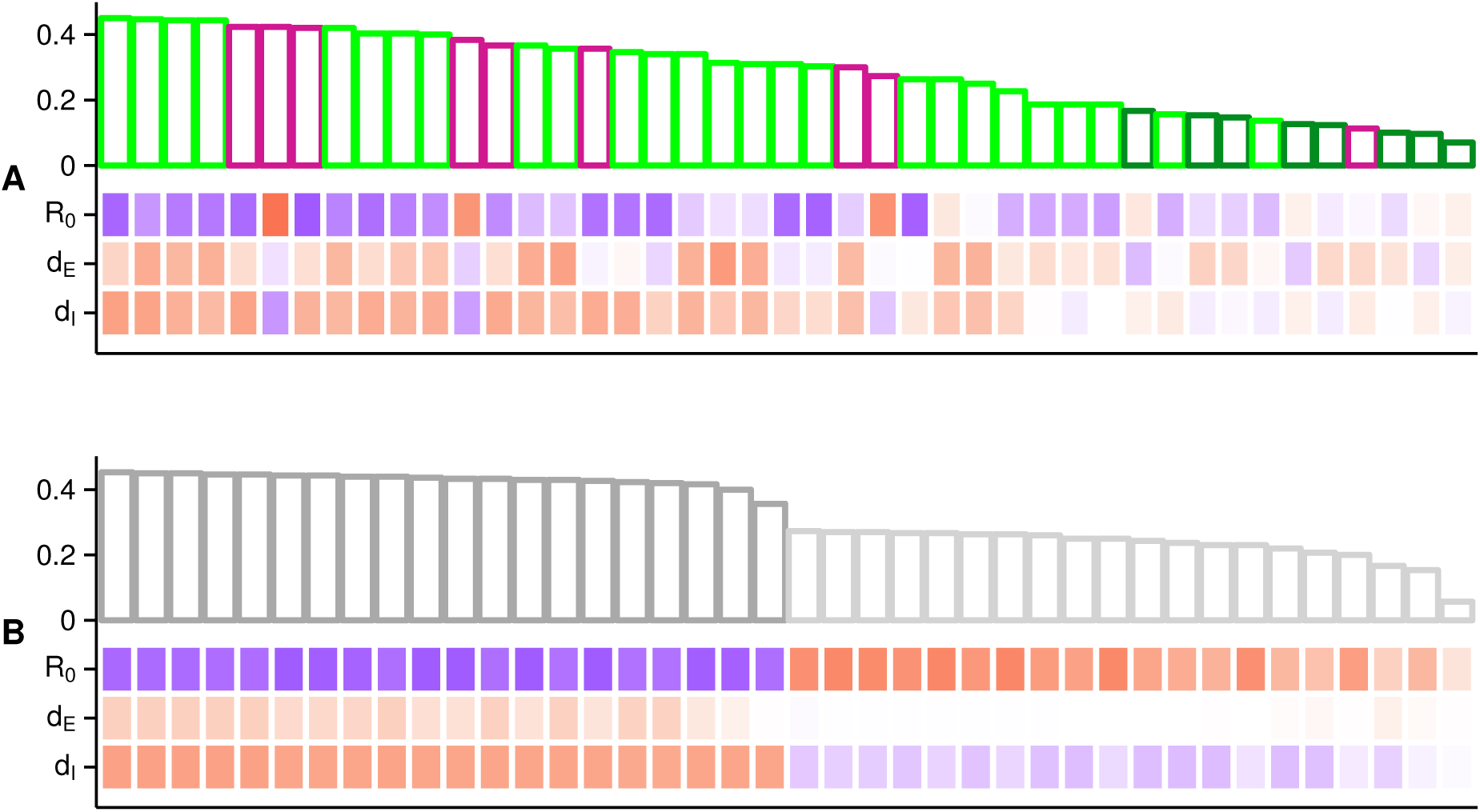
Heat map and histograms of Spearman’s correlations between epidemiological parameters of the BDEI model and all sets of summary statistics for trees of 72 leaves simulated assuming p ≈ 0.4. In panel A, the colors correspond to the bl (light green), topo (dark green) and ltt (magenta) sets. Panel B show the coords set related to the LTT plot with x-axis (dark gray) and y-axis (light gray) coordinates. On x-axis, summary statistics or coordinates are ranked from the most to the least correlated to all epidemiological parameters. Bar heights in the histograms represent the mean absolute correlation of each summary statistics to the whole set of parameters. Summary statistics and coordinates are ranked from the most to the least correlated. Correlation values between each summary statistics (or coordinate) and each epidemiological parameter are displayed in the heat map, where squares are colored according to a gradient from red (highly positively correlated) to white (no correlation) and blue (highly negatively correlated).

In this context of data analysis, it is important to assess the fitness of the summary statistics to infer the epidemiological parameters from the ‘target’ phylogeny. We did this for the sumstats and coords sets together and separately. The goodness-of-fit test revealed that the coords set of summary statistics is not fit to infer the epidemiological parameters of the Ebola phylogeny (p-value of the goodness-of-fit test lower than 0.05). Therefore we inferred parameters from the Ebola phylogeny using only the sumstats set of summary statistics.

Fig. 9 shows that the median of the posterior distribution of *R*_0_ inferred by Stadler et al. using BEAST2-BDEI, is close to the median of their prior distribution (in gray). The duration of latency seems very difficult to infer using the BEAST2 approach, because the *d*_*E*_ HPD 95% is almost as large as that of the prior.

**Figure 9.**
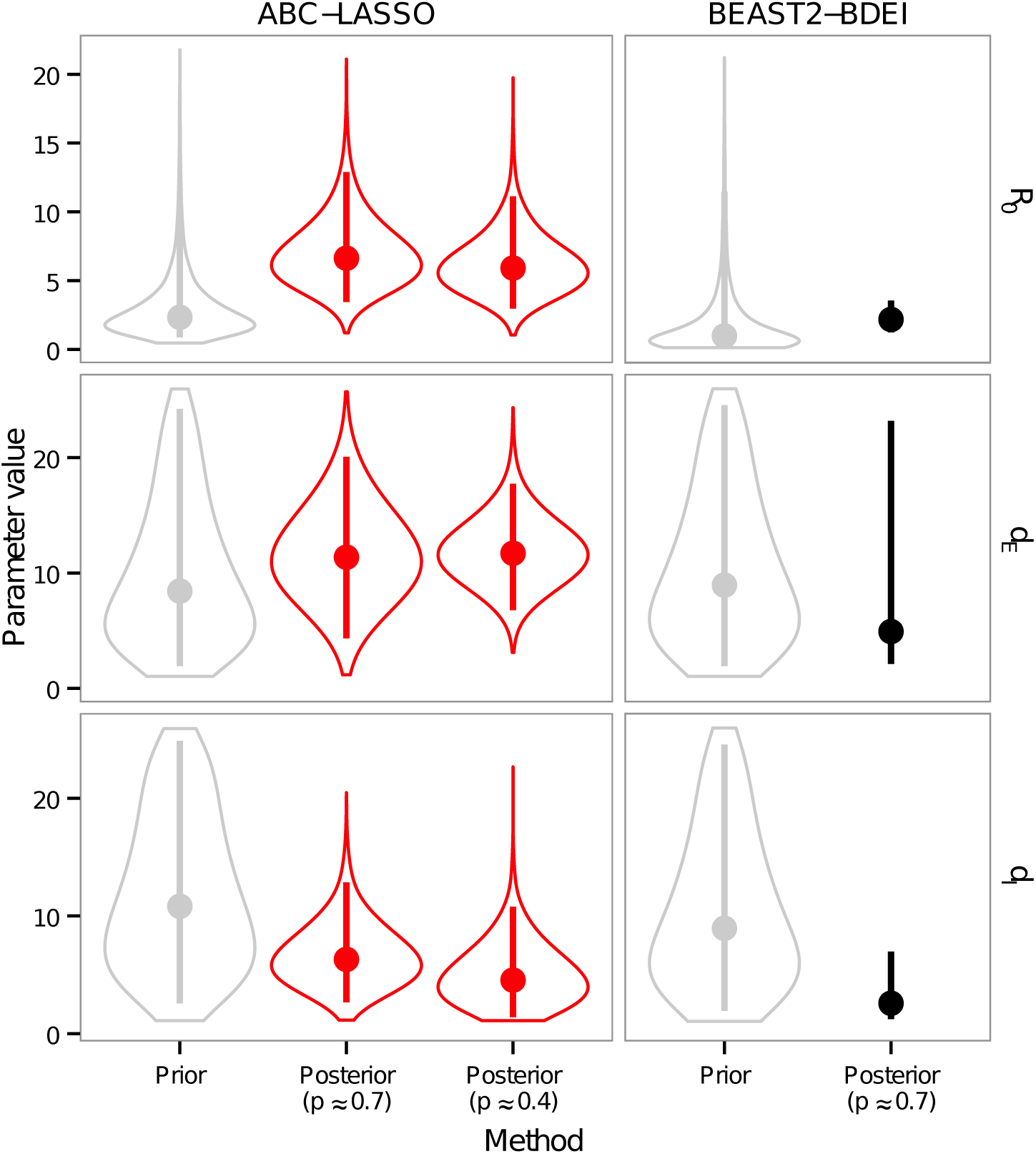
Prior and posterior distributions of parameter estimations from the Ebola phylogeny. We show the results for three different inference methods: ABC-LASSO (in red) and BEAST2-BDEI from Stadler et al. (in black). Gray violins correspond to the prior distributions and red violins to ABC-LASSO posterior distributions. The dots represent the median and the vertical lines the 95% highest posterior density of each distribution.

Our parameter estimates slightly differ from Stadler et al.’s but they are biologically relevant according to results obtained using different epidemiological methods. We find a longer incubation period (11.7 [HPD_95%_: 6.77 - 17.74]) and a longer duration of infectiousness (4.5 [HPD_95%_: 1.41 - 10.79]) than Stadler et al’s (4.92 [HPD_95%_: 2.11 - 23.20] and 2.58 [HPD_95%_: 1.24 - 6.98] respectively). Both of these are more in line with the estimations from the WHO Ebola Response Team [65], which found that the fitted incubation period was 9.9 ± 5.6 days and the mean duration of infectiousness in the community was about 4.6 ± 5.1 days. We also infer a greater value for the *R*_0_ than Stadler et al (5.92 [HPD_95%_: 2.97 - 11.12] instead of 2.18 [HPD_95%_: 1.24 - 3.55]), which is probably driven by the longer duration of latency. Indeed, even if the duration of latency does not appear in the deterministic formulation of the *R*_0_ for the BDEI model, in the stochastic setting it may have an effect. Put differently, in our simulations, we have more infected individuals but a high proportion of these individuals are still latent and do not propagate the disease.

## Discussion

Extracting epidemiological information from pathogen phylogenies largely remains an open challenge, especially for large phylogenies and complex models [13]. Here, we show that an Approximate Bayesian Computation (ABC) approach based on a large number of summary statistics to describe the phylogeny offers a promising alternative to existing methods.

There are two ways of performing ABC: either by using statistics to first summarize the data and then a simple distance function to compare the summary statistics values, or by using a more sophisticated distance function (sometimes called ‘functional distance’) to directly compare two datasets (observed and simulated). Summary statistics are sometimes viewed as ABC’s Achilles’ heel because the action of summarising suggests a loss of information. Furthermore, complex objects such as phylogenies can contain information that is not related to epidemiological parameters, which may dilute the inference signal. For instance, our results show that the topological shape of the phylogeny does not contain much information about the epidemiological parameters of both the BD and the SIR models, compared to the LTT plot and the branch lengths. This suggests that selecting only the ‘relevant’ part of the information in the phylogeny using summary statistics, could help improve the estimates of the epidemiological parameters. The problem is that selecting a few good statistics is notoriously difficult [41,66–68]. Here, we show that recent machine learning techniques can be used to perform variable selection on a large number of summary statistics, thus concentrating the inference signal. Removing the delicate step of choosing a subset of summary statistics opens wide perspectives for ABC.

In our ABC approach, we compute Euclidean distances between vectors of 83 unweighted summary statistics, some of which are highly correlated. We did consider weighting these summary statistics before calculating distances. The problem is that coming up with such a weight is not trivial given that the importance of each summary statistic for the inference can be affected by the epidemiological model or scenario considered. This is illustrated by the LASSO regression. This method efficiently performs variable selection in our ABC approach but when analysing the regression models, we were unable to identify sets of summary statistics that were always selected or always discarded. Adaptive methods of distance weighting exist but they are time consuming and tend to be replaced by sophisticated machine learning techniques. Finally, improving the rejection step might not be necessary since we use machine learning techniques to subsequently adjust the posterior distribution.

We compared two regression methods and concluded that regression using LASSO should be preferred to FFNN when using numerous summary statistics that are potentially correlated. This conclusion was largely driven by the fact that ABC-LASSO was more robust to summary statistics choice than ABC-FFNN. However, this is highly dependent on the R packages used (glmnet and nnet) and we expect that a re-implementation of a FFNN model with regularization tuning could give results at least comparable to that of the ABC-LASSO.

Another possibility for the regression step could be to use random forest algorithms, which are powerful tools for clustering and non-linear regression [69]. The supervised classification algorithm has already been used in an ABC context [70], and the regression algorithm could be used as a non-linear model for posterior adjustment. There could be two advantages in using the random forests regression algorithm instead of FFNN: first, as with LASSO, variable selection can readily be quickly optimized with random forests and, second, it provides information about the retained variables and the resulting regression model, while FFNN is a ‘black box’.

Using robust machine learning techniques that perform non-linear regressions, questions the usefulness of the initial rejection step. Indeed, why discard data if we can perform a non-linear regression using variable selection? In fact, with those machine learning techniques, the larger the learning set size, the more accurate the regression model. This is consistent with our result that ABC-FFNN results are more accurate when the tolerance threshold is high (i.e. when rejection is minimal). Overall, new non-linear regression techniques may challenge the canonical rejection step of the ABC, but this is beyond the scope of this study.

We compared our approach to methods implemented in the framework package BEAST2 because they are based on the expression of the likelihood function, meaning that they are robust methods, and also because they are popular and accessible. Several other interesting methods exist, either based on explicit likelihood functions (e.g. coalescent approaches [20]) or ABC approaches (e.g. using a functional distance [42]). Comparing all these methods would of course be valuable but is challenging because not all of them have been released as stand alone packages.

Interestingly, we show that comparing posterior distributions is valuable because statistics such as the relative error are not sufficient to evaluate the performance of an inference method and may lead to error-prone conclusions. For instance, if the prior distribution is approximately centered on the targeted value, without any selection on parameter values the posterior will not deviate from it. In fact, for the SIR model, the relative error suggested that ABC-FFNN outperformed ABC-LASSO if only some of the summary statistics were used (see Fig. 5). However, when we examined the posterior distributions, we found that ABC-LASSO posterior distributions had deviated further from the prior distribution than ABC-FFNN distributions (results not shown). More generally, the shape of the posterior distribution is extremely informative about the goodness of the fit.

One of the major challenges in phylodynamics has been to extend simple epidemiological models to more complex and realistic systems. However, many methodologies seem to rapidly reach their limits due to a trade-off between model complexity and computation time. This trade-off becomes even more acute as the phylogeny size increases. In this study, we show that ABC approaches are less limited by model complexity and phylogeny size. Moreover, more complex models could be easily tested using ABC-LASSO, since the major requirement of our approach is to be able to rapidly simulate data assuming such models. Indeed, we chose summary statistics that are computable in linear-time complexity, so that their computation take little time compared to simulation.

We show that topological statistics contain little information about the epidemiological parameters of BD and SIR models. However recent studies reveal that these statistics may become useful for parameter inference when considering more complex models such as models including spatial structure [41] or risk structure [42, 71]. It would therefore be interesting to re-analyse these models with our new approach.

With the constant decreasing cost of sequencing technologies, epidemiological studies of viruses provide larger phylogenies and we need fast and effective methods to analyse them. Current methods all tend to reach their limits for simple non-trivial models (e.g. the SIR model) when the size of the phylogeny increases. ABC approaches involving many summary statistics and a regression step offer a promising and flexible alternative. Not only do they allow to optimize the choice of summary statistics, but also their computing time seems to be less limited by phylogeny size and model complexity than existing likelihood-based methods.

Finally, we have focused here on phylogenies of infections but this method could be extended to infer parameters from phylogenies generated using ecological or evolutionary models [72, 73].

## Acknowledgments

The authors thank Carmen Lía Murall, Mircea Sofonea and Louis Du Plessis for useful comments on this research. ES is grateful to Vincent Lefort for the technical support.

